# Improved bacteria population structure analysis on thousands of genomes using unsupervised methods

**DOI:** 10.1101/599944

**Authors:** Katrina Schlum, Se-Ran Jun, Zulema Udaondo, David W. Ussery, Scott J. Emrich

## Abstract

Over ten thousand genomes of *Escherichia coli* are now available, and this number will continue to grow for this and other important microbial species. The first approach often used to better understand microbes is phylogenetic group analysis followed by pan-genome analysis of highly related genomes. Here, we combine sequence-based features with unsupervised clustering on up to 2,231 *E. coli* genomes and a total of 1,367 *Clostridium difficile* genomes. We show that Non-negative Matrix Factorization (NMF) can identify “mixed”/cryptic genomes, and can better determine inter-related genome groups and their distinguishing features (genes) relative to prior methods.

## I. Introduction

The advent of inexpensive whole genome sequencing has led to the generation of thousands of related genomes, even from a single study (e.g., [1]). Large-scale genome analysis of *E. coli* is of particular interest given its importance in experimental, biotechnology and health-focused research. Systems biology applications, for example, could benefit from a “streamlined” chassis genome containing only essential genes. Finding such essential genes, however, is complicated by functional redundancy that can compensate for individual gene knock outs [2], while other genes may only be required only under certain conditions/habitats.

Computational analysis of deeply sequenced species initially focused on two applications: defining a “pan-genome,” which is the set of all observed sequences across all samples, and defining a “core genome,” the relatively stable subset found in nearly all of its members (see [3], [4] for reviews). In *E. coli*, prior computational analysis using different genomic markers has also shown that this species can be divided into five distinct interrelated groups, or phylogroups, denoted A, B1, B2, D or E. A sixth group consisting of named *Shigella* species is somewhat contentious as to its placement(s), as is a potential seventh ancestral group called “Clade 1” [1]. This characterization is also likely incomplete since a group F has been recently reported, as well as a another new group C based on PCR-based sequence variation [5].

Here, we use data-driven computational methods to predict interrelated groups within a single bacterial species. First, we confirm that sketching based on a random sample of k-mers (see [6], [7]) can uncover variation consistent with the known *E. coli* phylogroups. We then focus on unsupervised clustering and specifically Non-negative Matrix Factorization (NMF; [8]). NMF has been rediscovered for clustering metagenomic samples [9], [10] and, more recently, to cluster and refine marker genes for single cell RNAseq [11]. In this study we consider a new application: analyzing gene presence/absence to reveal population structure within two bacterial species.

Our main contributions are a new approach to determine atypical bacterial genomes relative to known (and inferred) groups and show examples of substructure within existing phylogroups that are not currently captured by traditional multilocus sequence typing (MLST) profiles [5]. We show that NMF can augment traditional large-scale computational analysis of genome collections, especially when labeled genome sequences are available.

## II. Methods

### A. Data used in this study

Three published collections were used: 2,244 *E. coli* whole genomes [1], 186 *E. coli/Shigella* whole genomes [4] and 59 *Shigella* whole genomes [12]. All genome assemblies were downloaded from GenBank and the NCBI Nucleotide database on May 10, 2018. The fourth unpublished dataset used was all 1,367 *Clostridium difficile* complete genomes available from GenBank as of August 22, 2018. For all datasets, plasmids and multiple chromosomes were included as one sequence file. All pan-genome family matrices were generated using USEARCH [13] where each gene family member has a least 50% sequence identity and length coverage of at least 50% of the query sequence length using an in house script as previously described in [14]. To demonstrate this method on a larger dataset, a collection of 2,231 *E. coli* genomes was used from a Genome Announcement set [15].

### B. Determining groups using sketches (Mash)

A number of groups have shown that MinHash sketching, a technique first invented to detect duplicate web pages for Alta Vista [16], is an effective estimate of genome distance [6], [7]. Briefly, MinHash sketching uses a fixed vector of random hash values per sample to estimate the genomes’ k-mer based Jaccard similarity (distance). The primary advantages are computational; each “sketch” compresses a *∼*5.0 MB *E. coli* genome into a *∼*100k file, an immediate 50X improvement in space and a substantial improvement in runtime given that sketches are directly comparable [16]. Sketching can also be applied to unassembled data (see [6] for *E. coli* comparisons using different sequencing platforms).

Here Mash [6] was used to represent each genome as a MinHash sketch with 10,000 random hash functions (21 k-mers), the value of which was empirically determined for *E. coli* in [7], followed by calculating the distance between each sketch using Mash. Although four different hierarchical clustering methods (UPGMA, Wards, complete, average) were initially applied to the resulting matrix, the *hclust* complete function in R was deemed the best and used throughout (data not shown). Clusters were visualized in R as dendrograms using R’s *dendextend* [17] and *plot*.

### C. Unsupervised clustering

Principal component analysis (PCA) is traditionally used to reduce data dimensions, visualize similarity, and filter noise. We applied PCA using the Pandas Python package [18] to the pan-genome family count matrix with whitening. Initial runs indicated large and relatively unique signal primarily in *Shigella* genomes thus a log base 10 + 1 scaling transformation was applied. Non-negative matrix factorization (NMF) was also used to help identify the relevant gene families that defined the interrelated groups. NMF was first introduced in 1999 by Lee and Seung to reduce large image bit maps and identify important features [19]. NMF can identify important patterns that are driving resulting clusters and assign membership using the index of the dominant basis component of samples (columns) or features (rows). Our analysis used the NMF R package [20] with an initial cluster size of six based on the previously described groups A, B1, B2, D, E, and *Shigella*. NMF results were visualized in R using pheatmap [21].

### D. Supervised clustering

Random forest (RF) classifiers are a type of divide and conquer ensemble where a set of decision trees are generated to create rules based on identifying features and targets from a training data set. The votes for each decision tree are aggregated and a majority rule is used to predict the class from the test data set. We used the pan-genome family count matrix as input to the RF classifier, which was trained using 1000 decision trees with 80% of the data set and tested on 20%. The RF classifier used here was from scikit-learn [22].

## III. Initial Results

### A. Mash vs. Core genome-based clusters

We first assessed sketch-based classification relative to data from [4] for which traditional phylogenetic group analysis was performed: we were able to similarly classify all but twelve genomes (see Figure 1 for a broad visualization) and were better able to detect Clade 1 relative to [4].

**Fig. 1.**
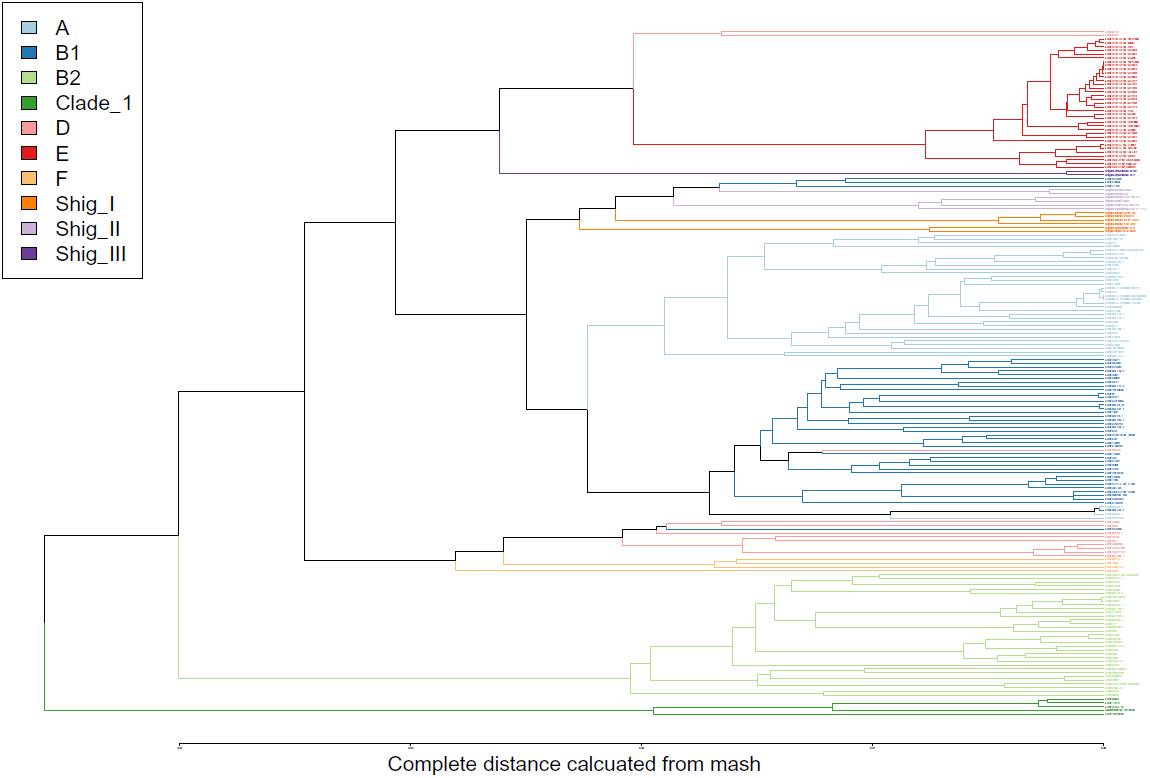
Mash Tree based on 186 genomes [4]

Some of the overall disagreement with [4] may result from a small sample size and/or relatively lower quality draft genomes not well suited for sketching. To rule out data size concerns, the twelve disagreement genomes from [4] were added to a larger 2,224 *E. coli* dataset [1]. Only 33% (3 genomes) of the Mash disagreements were now placed properly; Mash, ribosomal assignments (data not shown), and a third method— FastANI-based clustering [23]—all agreed on the remaining nine placements. ez-Clermont, a computational platform based on established diagnostic genes, can independently assess existing labels, even though it did not classify as well (24 total inconsistencies). Seven ez-Clermont labels agreed with the prior three methods, with the remaining two failing to provide any label. Interestingly, sketch-based results also uncovered Clermont’s novel “C” group [5] in the same relative location (between B1 and D; see Figure 1).

Another likely source of assignment difficulties are the *Shigella* subgroups [24] and pathogenic *Shigella*-like *E. coli*, especially since *Shigella* was under sampled in [4]. To address this we next added 59 *Shigella* genomes from [12] to those from [4]. Although all non-*Shigella* genomes clustered as before, two prior disagreements were now clearly *Shigella*. These results suggest the sensitivity of *E. coli* and *Shigella* sketches slightly improve when provided more data, as expected.

### B. Traditional clustering based on “pan-genome” data

The pan-genome in *E. coli* is predicted to be an “open” one (infinite) [4], and is 30,225 gene coding sequences in our “traditional” 186 *E. coli* pan-genome dataset with no singletons, i.e., genes found in only one genome. The pangenome family matrix of the larger 2,224 *E. coli* genome set was 60,654, also with no singletons, and computational analysis of gene presence/absence has often been used to infer *E. coli* interrelated groups in part since certain genes/pathways may be group-specific (see [4]).

Such genome content similarities can be visualized using Principal Components Analysis (PCA). For the genome data from [4], the first component captures diversity in non-E genomes, the second principal component separates E genomes from others, while the third component isolates *Shigella* vs. non-*Shigella* (data not shown, but see Figure 2 for similar results). After using filtering similar to [4], *Shigella* samples (and esp. *Shigella* III) are the most distinct, followed by the B2s. Clade 1 genomes, which are presumed ancestral, were found on the axis of B1, E and A, as expected [4]. Given gene copy number differences largely in *Shigella*, we applied a log + 1 transformation (Methods) that appears to better highlight non-*Shigella* genomes, esp. the genomes belonging to the new F group and between D and B2 (Figure 2). Although this PCA ordination intermixes the closely related B1 and A genomes, as well as Clade 1 with its relatives, it supports a D-like assignment of disagreeing GCA 000194235.2, similar to the assignment from all prior methods.

**Fig. 2.**
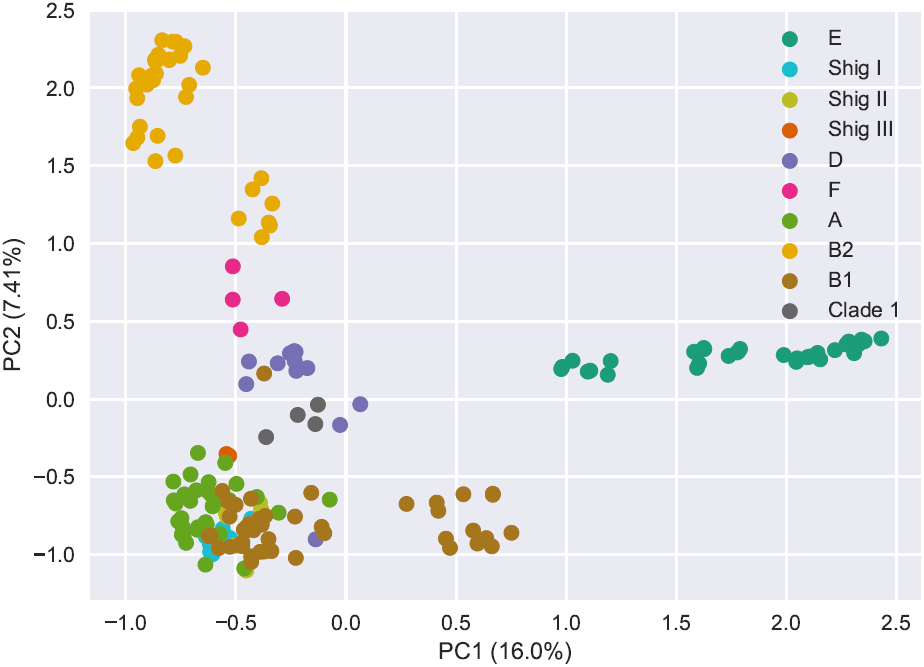
PCA ordination on gene family count matrix of 186 genomes [4]

We next performed random forest (RF) analysis since RFs have been previously effective in multiple microbial applications (see [25]). Similar to the Mash-based clustering, overall accuracy was high 0.999 (+/-0.001) for the 2,244 Genome Announcement dataset and relatively lower for the 186 genome set (0.939 +/-0.23). The confusion matrices were plotted and no false positives or negatives for the 2,224 were clearly visible; however, there was ambiguity between Clade 1, D and F on the smaller number of genomes (Figure 3), just like in the PCA ordinations in Figure 2.

**Fig. 3.**
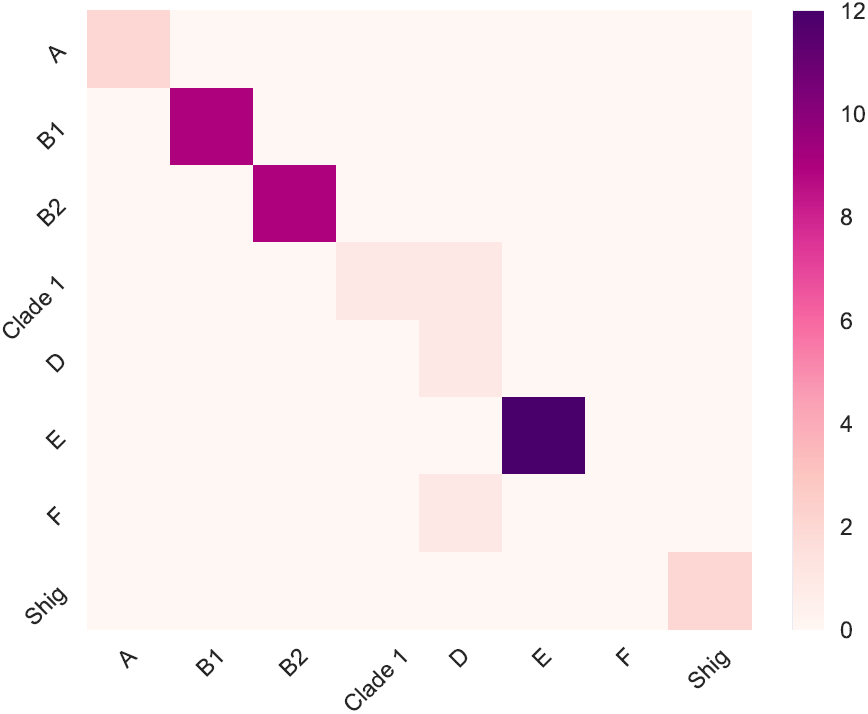
Random forest confusion matrix on gene family count matrix of 186 genomes [4]

**Fig. 4.**
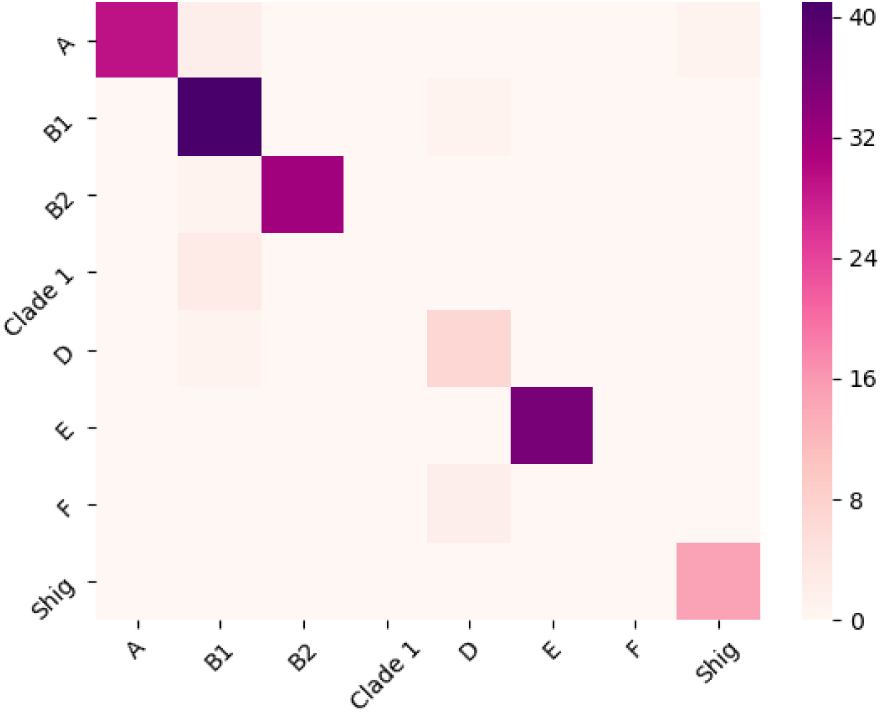
NMF confusion matrix (k = 10) based on predicted labels in [4] with bootstrap value *>*=0.6 on 10 NMF runs

In summary, a total of four genomes were consistently in disagreement using the core genome alignment group assignments relative to six independent data-driven methods considered, which all agreed on their placements. Based on these results we conclude gene presence/absence data can be used on thousands of genomes to understand most interrelated groups, but like genome sketches these methods can be limited when applied to smaller collections of genomes.

## IV. Using NMF to predict interrelated groups and atypical genomes

Every computational approach considered so far does has not worked as well on less defined subgroups such as *E. coli* Clade 1. NMF [8] can partition a set of features into a fixed number of components/groups. Further, because of random initialization used by NMF, its results can vary between runs. To account for and to leverage this variation, we perform bootstrapping similar to phylogenetic tree analysis to better identify and hopefully characterize atypical bacterial genome content using a gene presence/absence matrix as input.

When first applied to the 186 *E. coli* genome pan matrix, with six pre-defined subgroups, NMF consistently partitioned the following: all E (n=36), the majority of the *Shigella* (n=13), B2 (n=30), A (n=32), B1(n=30), D(n=11), consistent with the RF confusion matrix in Figure 3. Clade 1 members, however, were still found to be ambiguous.

We next performed 10 NMF runs with 6 groups and measured assignment stability. Interestingly, genomes with a bootstrap value of 1.0, i.e., were always assigned to the same group across runs, were in complete agreement with the established phylogroup labels with Shigella I/II as a single group. 20/25 of the genomes with a confidence of 0.9 were consistent: two F genomes were labeled as B2; GCA 000188815.2 was labeled as a Shig, matching the same placement as Mash; GCA 000167915.2 that appears on the edge of A/Shigella was labeled as *Shigella*, and the Mash outlier GCA 000194215.2 was placed in E. There were 10/13 consistent genomes with a confidence of 0.8 where all three inconsistent genomes encode for at least one Shiga toxin. When we used a 0.7 cutoff, there were a total of ten genomes that were deemed atypical, i.e., inconsistent with a single group. In total, NMF labeled four new genomes as inconsistent relative to Mash (2 F’s and 2 A’s). Of the twelve Mash-based inconsistencies, 5 were potentially resolved by NMF and as a result NMF performs slightly better in terms of matching established *E. coli* phylogroup labels. Significantly, NMF agreed with Mash (and other methods) on most of the atypical genomes relative to [4] (7/12 genomes). Of the remaining five, 60% (3 genomes) had bootstrap values *<*0.6, indicating NMF can assign lower label confidence for these genomes.

A substantial advantage of NMF is that it will best assign structure using any number of pre-defined groups, unlike the other methods considered here. For comparison, and to potentially identify novel group substructure, we ran NMF with a number of group counts and found k = 10 was the best on these data; there was 100% consistency between 87 genome’s group assignment with a confidence of 1.0 and with the 40 genomes assigned confidence of 0.9. Separation of the groups was also clearly visible using a heatmap of the NMF consensus matrix (Figure 5). Additionally, 14/15 genomes were consistent when the confidence threshold was decreased to 0.8; the one inconsistent genome as D (GCA 000194435.2) was always classified by other methods (inc. NMF with k=6) as B1. This genome is also a pathogenic *E. coli* with stx1 and stx2. So by using ten groups and a bootstrap cutoff of 0.8 consistency reaches as high as 100% on a total of 142 genomes, while consistency on the same data is 94% with NMF (k=6) and also with random forest classification.

**Fig. 5.**
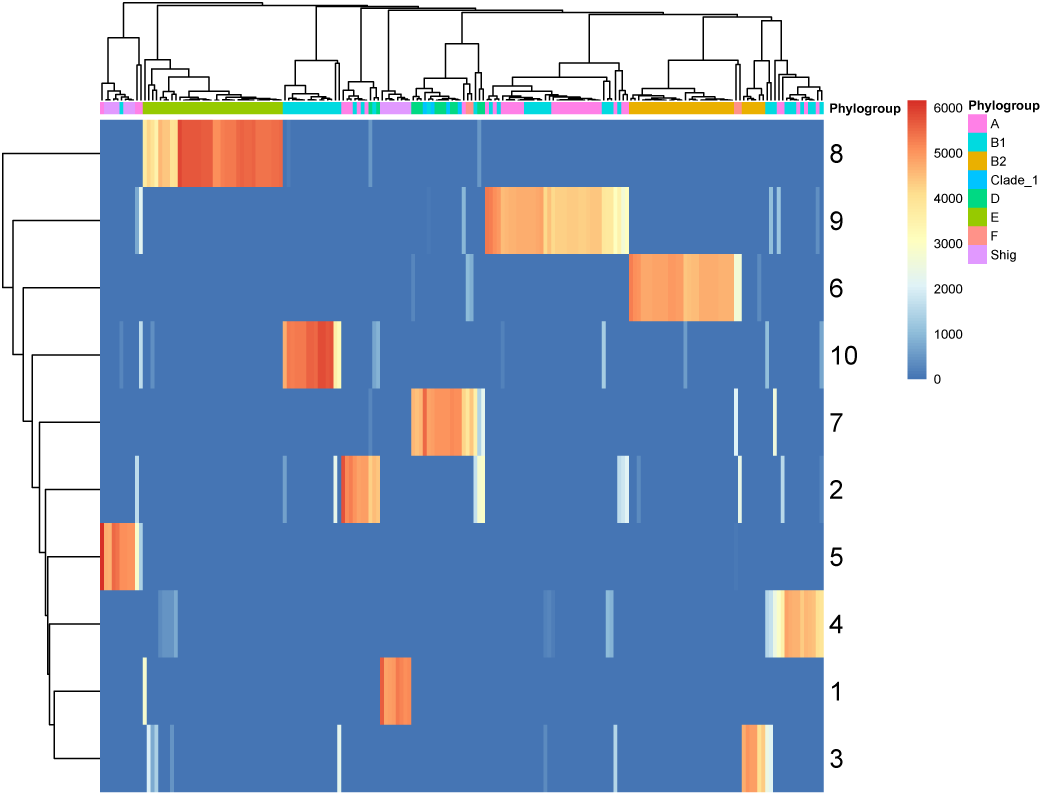
Consensus matrix computed from one NMF run with k = 10

A total of 24 genomes had a confidence *<*0.6 with k=6 including all four “Clade 1” (D as predicted by NMF) and twelve genomes currently labeled as D. We note that only seven of these low confidence genomes previously showed a conflicting assignment. We speculate that NMF may be providing higher resolution, especially since there were only eleven total incongruent genomes when k = 10. NMF with ten groups also had higher confidence: fewer genomes had bootstrap scores less than 0.7 (14 genomes) and it is clear that, for example, NMF with 10 groups can better identify the D phylogroup (4 vs. 11 inconsistencies) relative to the core genome alignment “gold standard”. To support the increased accuracy observed with k = 10, we plotted the cophenetic correlation coefficient and residual sum of squares (RSS). The cophenetic correlation coefficient shows the correlation between the dissimilarities and the cophenetic distance of each pair of observations. For the cophenetic, we observed a dip point at 10 which suggests 11 as largest number of possible groups (data not shown). The RSS defined as the sum of the squares of the residuals (deviation of predicted from actual) showed an inflection point at k = 9 with the RSS plot (data also not shown); both are greater than the six primary groups currently thought to exist for *E. coli*.

To evaluate the number of potential phylogroups beyond six, we plotted the phylogroup splits on one NMF run using the 186 *E. coli* dataset as illustrated in Table I. Two stable trends were observed as the number of groups (k) was increased: B1 always split at least 2 times and E was always split twice. We propose that NMF may be detecting associations (subgroups/interrelationships/ecological similarities) that are not as clear when using the genome-wide similarity methods considered previously.

**TABLE I.** PHYLOGROUP SPLIT ON NMF RUNS FOR 186 GENOMES

| K | A | B1 | B2 | D | E | Shig |
| --- | --- | --- | --- | --- | --- | --- |
| 6 | 1 | 1 | 1 | 1 | 1 | 1 |
| 7 | 1 | 2 | 1 | 1 | 1 | 1 |
| 8 | 1 | 2 | 1 | 0 | 2 | 2 |
| 9 | 1 | 3 | 1 | 1 | 2 | 1 |
| 10 | 1 | 3 | 1 | 1 | 2 | 2 |

To further validate our NMF framework, we used the same large 2,231 *E.coli* genome collection, using 10 bootstrap runs each, on k values ranging from 6 to 10. We achieved 95.1% accuracy with 0.7 as the bootstrap value (k = 9) where 80.4% of the 163 atypical genomes were E/D and all 45 of the F genomes were classified as B1/B2/D. These results suggest that phylogroup D/F/E genomes are the most difficult to consistently classify and the A, B1, and B2 groups are more established and/or stable, at least in this collection.

### A. Applying NMF based classification to Clostridium Difficile genomes

We also processed 1,367 *C. diff* genomes through our NMF framework since biologists do not believe this species has the same latent genome content structure as well-studied *E. coli* [26]. The pan-genome family count matrix for this dataset had 55,186 singletons that were filtered prior to analysis, resulting in a 1,367 (genomes) x 8,871 (non-singleton genes) matrix. MLST labels were generated from the software mlst developed by Trosten Seemann that imports *C. diff*’s MLST schema from PubMLST [27], [28]. Since *C. diff* is a more diverse species compared to *E. coli*, the high number of singletons found was expected in the pan-genome calculations.

The population structure of *C. diff* is thought to include 8 clades based on the most recent core genome MLST assignments [29] but the current PubMLST only includes profiles for 5 of these groups. Based on the current PubMLST profiles, the 1,367 *C. diff* genomes analyzed were identified as phylogroup 1 (n=870), phylogroup 2 (n=239), phylogroup 3 (n=14), phylogroup 4 (n=73), phylogroup 5 (n=55), and 114 were untypeable. Additionally, 5 groups was supported by the inflection point on the cophenetic correlation plot. Therefore, we first ran NMF with k = 5 and observed a consistency of 93.7% (1032/1110) using a bootstrap cut-off of *>*= 7/10 and excluding the 114 untypeable genomes. We observed that clade 3 always grouped with clade 1, clade 4 always grouped with clade 1 with two exceptions, and clade 5 always grouped with clade 2; however, this genomic dataset is highly biased towards clade 1 and 2.

To determine the effect of the observed sample bias on clade 4 and 5 assignments, we randomly down-sampled clade 1 to a total of 239 genomes, removed clade 3 and another four contaminated genomes marked as none type per MLST and identified as most similar to Eggerthella and Terrisporobacter rather than *C. diff* per Mash comparisons to the Refseq database [6]. To evaluate the robustness of NMF assigned phylogroup labels using this down-sampled dataset of 716 genomes, we applied bootstrapping to our NMF label assignments on 10 NMF iterations (k=4). Although we were unable to type 110 genomes (“none”) for comparison, this revised analysis resulted in all clade 4 genomes grouping together and likewise for clade 5 with an overall consistency of 95.2% (556/584) using a bootstrap cut-off of *>*= 7/10. Only 32 genomes had a bootstrap value *<*7/10; these genomes may belong to three novel MLST groups that are not currently accounted for in the current MLST schema or be more atypical. A total of ten genomes were labeled as none per pubMLST but were assignable by NMF as either clade 1, 2 and 5 (bootstrap cut-off of *>*= 7/10). This indicates NMF is able to handle untypeable genomes unlike pubMLST, which is reliant on *a priori* knowledge present in a database.

We also observed increased detection capabilities on the full *C. diff* dataset. From the 114 pubMLST untypeable genomes, our NMF framework was able to assign a total of 87 genomes to either clade 1, 9 or 2 (bootstrap *>*= 7/10, k=5). This suggests that the MLST pre-defined groupings may not be capturing the allelic profiles for at least 87 of these genomes but our method is able to label them based on global genomic content similarities. Interestingly, when NMF is ran with k = 8, 74 of the atypical *C. diff* genomes (bootstrap of 0.6 or lower) appear to be a mixture of four clades, which suggest these some of the *C. diff* genomes are more heterogeneous than others and may never be classified into one phylogroup. We note that it is possible that these “mixed” genomes may ultimately also be assigned to MLST based clade I, which has previously also characterized as the most heterogeneous *C. diff* clade [30].

## V. Discussion

We obtain similar accuracy using two completely different bacterial genome collections without having prior group labels (either based on a core genome analysis or MLST). These results suggest that the application of NMF to pan-genome matrices is a more sensitive approach to determine interrelated genomes, and with higher confidence via bootstrapping. We find that high bootstrap values correspond better to known groups, esp. when a higher value is provided to NMF: 10 for *E. coli* and 8 for *C. difficle*.

Prior methods define genomes on a continuum, say using PCA ordination. Our NMF-driven method uncovers associations based on shared gene content. In support of this, we find one *Shigella* cluster with NMF but up to three separate clusters based on genome similarity. Because both the number of groups and the bootstrap value are flexible, it is possible to be more or less conservative as desired.

Unlike alternative tree/clustering-based methods that explain additional groups via subclades/groups (e.g., C and F), it is unclear what drives the 10 inferred gene-based *E. coli* groups versus the traditional six. We hypothesize that the data-driven NMF approach is uncovering additional “cryptic” structure even in well-studied *E. coli*. Interestingly, many of the inconsistent assignments within *E. coli* encode at least one toxin, and we also note that both outgroups (GCA 000176575.2 and GCA 000163175.1) have low confidence in NMF sample probability assignments.

Further, although its not as obvious how to apply the terms “introgression” and “admixture” to bacteria, we believe NMF may be able to determine atypical genomes that have absorbed and/or still contain non-traditional features (genes). For example, *E.coli* “Clade 1” is difficult to assign due to appearing to have an even mixed of features that are better associated with other defined groups present in these data.

## VI. Conclusion

Based on our approaches for classifying bacterial genomes, we have predicted that at least seven public *E. coli* genome instances may be “mixed,” i.e., includes genes more typical of other interrelated groups. To the best of our knowledge we have shown the first means to uncover these atypical genomes. For example, these seven genomes are inconsistently classified in both NMF and Mash-based distance while PCA ordination supports at least three of these genomes as a mixture.

We show that the six historical *E. coli* interrelated groups can also be defined by the presence/absence of gene families with an accuracy of 93.5% (bootstrap *>*= 0.7) using NMF with k set to 6 and up to 100% accuracy (bootstrap *>*= 0.8) when 10 groups are set. We also show similar accuracy and results on a completely different bacteria species, *C. diff*, for which biologists believe is less structured than *E. coli.*

NMF is a new way to further our understanding of bacteria genomes. Based on feature importance, we can next predict phylogroup specific genes, or “phylocore” genes, that consistently define the NMF-based groups. Defining phylogroups based on genome presence/absence similarities provides us a means to identify subgroup associated genes and, for example, remove them before performing bacteria genome wide association studies (GWAS). We also believe that starting to define genes that are shared within these inter-related groups could also associate to certain ecological niches (e.g., host-specific vs “environmental” in *E. coli*) and, ultimately, increase our understanding of the fundamental and functional differences of bacterial inter-related groups at the genomic level.

## VII. Acknowledgments

We would like to thank David Molik and RJ Nowling for very helpful discussions related to NMF and PCA, respectively. We would also like to thank members of the Ussery lab, specifically Kaleb Abram for helping us run Clermont typing and Skylar Connor for additional quality checks.

